# A multiple ion-uptake phenotyping platform reveals shared mechanisms that affect nutrient uptake by maize roots

**DOI:** 10.1101/2020.06.15.153601

**Authors:** Marcus Griffiths, Sonali Roy, Haichao Guo, Anand Seethepalli, David Huhman, Yaxin Ge, Robert E. Sharp, Felix B. Fritschi, Larry M. York

## Abstract

Nutrient uptake is critical for crop growth and determined by root foraging in soil. Growth and branching of roots lead to effective root placement to acquire nutrients, but relatively less is known about absorption of nutrients at the root surface from the soil solution. This knowledge gap could be alleviated by understanding sources of genetic variation for short-term nutrient uptake on a root length basis. A new modular platform for high-throughput phenotyping of multiple ion uptake kinetics was designed to determine nutrient uptake rates in *Zea mays*. Using this system, uptake rates were characterized for the crop macronutrients nitrate, ammonium, potassium, phosphate and sulfate among the Nested Association Mapping (NAM) population founder lines. The data revealed that substantial genetic variation exists for multiple ion uptake rates in maize. Interestingly, specific nutrient uptake rates (nutrient uptake rate per length of root) were found to be both heritable and distinct from total uptake and plant size. The specific uptake rates of each nutrient were positively correlated with one another and with specific root respiration (root respiration rate per length of root), indicating that uptake is governed by shared mechanisms. We selected maize lines with high and low specific uptake rates and performed an RNA-seq analysis, which identified key regulatory components involved in nutrient uptake. The high-throughput multiple ion uptake kinetics pipeline will help further our understanding of nutrient uptake, parameterize holistic plant models, and identify breeding targets for crops with more efficient nutrient acquisition.

**Significance Statement:** Nutrient uptake is among the most limiting factors for plant growth and yet has not been used as a selection criterion in breeding. This is partly due to the lack of high-throughput phenotyping methods for measuring nutrient uptake. Here we describe a novel high-throughput phenotyping pipeline for quantification of multiple ion uptake rates. Using this new phenotyping system, our results demonstrate that specific ion uptake performance by maize plants is positively correlated among the macronutrients nitrogen, phosphorus, potassium and sulfur, and that substantial variation exists within a genetically diverse population. The findings reveal components of regulatory pathways possibly related with enhanced uptake, and confirm that nutrient uptake itself is a potential target for breeding of nutrient-efficient crops.

---

For plant growth and development, availability of the macronutrients including nitrogen (N), phosphorus (P), potassium (K) and sulfur (S) is critical. The availability of these macronutrients as ions in soil is often at limiting quantities for optimal plant growth (1, 2). In agriculture, chemical fertilizers are widely used to enrich soils and enhance crop productivity, but their usage adds a significant cost to production. Moreover, fertilizer use in agriculture is neither economically nor environmentally sustainable, with as little as 10-30% of applied fertilizer being captured by crop roots, and the remainder lost through leaching, erosion and as atmospheric emissions (3, 4). Understanding the genetic potential of plants for nutrient acquisition is important for developing nutrient-efficient crops (5, 6).

For a plant to acquire nutrients, the root system must perceive, grow to and intercept nutrients from the soil environment. Nutrient acquisition efficiency is defined as the amount of nutrient absorbed on a root cost basis (7, 8). There are two main processes that constitute nutrient acquisition efficiency: (1) root exploration for nutrients with modification of root growth and root system architecture, and (2) nutrient exploitation capacity of roots for taking up local nutrients (7, 9). In recent years, advances in root imaging and deep learning analysis approaches have shown great promise for root exploration trait-based crop selection (10–15). Multi-scale research linking environmental nutrient availability across time of plant development will be required to understand the functional processes and mechanisms plants employ for nutrient acquisition. Dissection of these complex interactions will provide new opportunities to improve sustainable crop production with more nutrient-efficient cultivars.

Nutrients are spatially and temporally heterogeneous in the soil and, therefore, plants have evolved to have high and low affinity transporters for uptake across nutrient concentration gradients (16–21). Ion-uptake kinetics studies have been instrumental in uncovering these distinct uptake systems in roots across many nutrients and plant species (22–24). Ion uptake kinetics research to date has demonstrated that species level variation exists for nutrient uptake rates on a per root basis (referred to as specific nutrient uptake rate) (25), with a few examples of genotypic variation within the same species (26–28). However, the research field is critically understudied as most phenotyping efforts rely on isotope accumulation, which is a low-throughput and destructive means of measuring uptake rates and ignores the interplay between nutrients (24, 29–31). Most studies focus on uptake measures of a single nutrient from simple solutions, however there are some examples of multiple nutrients from more complete solutions (26, 32). While total shoot nutrient content is sometimes assumed to be a proxy for root uptake capacity, so many factors influence overall plant growth and root exploration that nutrient content may be an unreliable indicator of specific uptake rate by roots (reviewed in (25)). Maximizing uptake kinetics is generally assumed to be beneficial for plant growth, however the energetic costs are likely substantial and therefore uniting the studies of uptake and metabolic cost is needed.

Here, we describe a modular phenotyping pipeline called ‘*RhizoFlux*’ for high-throughput phenotyping of multiple ion uptake kinetics in plants. Using this platform, we simultaneously quantified specific nutrient uptake rates (nutrient uptake rate per length of root) of nitrate, ammonium, phosphate, potassium and sulfate for each of the Nested Association Mapping (NAM) population founder lines in maize. We found that specific nutrient uptake rates were distinct from total uptake and plant size traits as an uptake efficiency related trait. In addition, we found that the specific nutrient uptake rates for each macronutrient were positively associated with one another, with notable genotypic preference for particular ions, and that specific root respiration rate (root respiration rate per length of root) was a significant contributor to specific nutrient uptake rate performance. All macronutrient specific uptake rates were highly heritable and, therefore, could be utilized as breeding targets for improved crop nutrient uptake.

## Results

### Development of a high-throughput multiple ion uptake kinetics pipeline

A plant phenotyping pipeline, *RhizoFlux*, was designed to phenotype in high-throughput multiple ion uptake performance for the macronutrients N, P, K and S simultaneously. To achieve this, a custom hydroponic growth and uptake experimental setup was designed and coupled with a data analysis workflow using R scripts (Fig. 1).

**Fig. 1.**
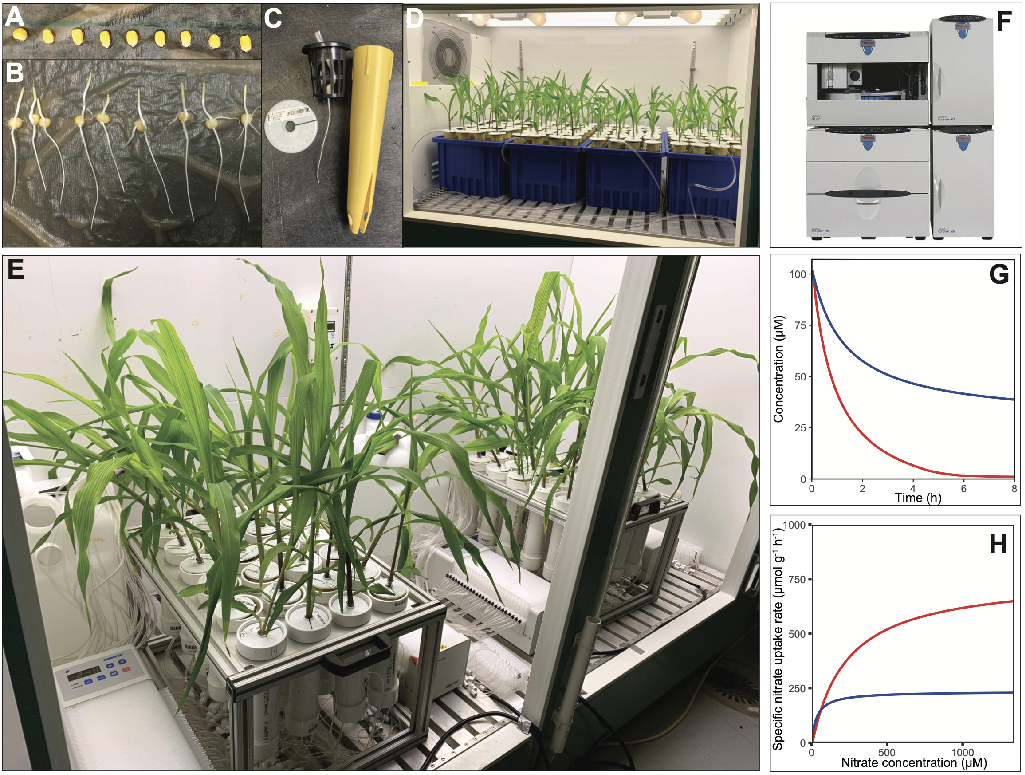
*RhizoFlux* platform for phenotyping of multiple ion uptake kinetics in plants. (A) Maize seeds are surface sterilized and germinated on germination paper rolls. (B) Evenly germinated seedlings are transferred to (C) seedling cones and grown in (D) aerated hydroponics for 14 days. (E) Plants are transferred to a custom ion uptake setup consisting of 24 hydroponic chambers connected to two 24-channel peristaltic pumps for simultaneous solution loading, aeration, and sampling onto a collection plate. (F) Ion concentrations of collected nutrient samples are determined using ion chromatography (G) for quantifying ion depletion across time and (H) used to calculate net specific ion uptake rates on a root length or mass basis.

To attain the experimental throughput and reproducibility necessary for mapping population-sized studies on plant nutrient uptake, two separate systems were designed, one for plant growth and one for uptake measures. Maize seeds were germinated, and seedlings grown together in large hydroponic tanks (Fig. 1*A–D*) and then transferred to individual hydroponic chambers for nutrient uptake phenotyping (Fig. 1*E*). This enabled the uptake measurements to be scaled up to greater numbers of lines with a time-staggered experimental block design. For measurement of multiple macronutrient uptake rates simultaneously, a basal nutrient solution was developed where the concentrations of calcium and 2-(N-morpholino)ethanesulfonic acid (MES) buffer were kept consistent to minimize pH fluctuation and preserve root membrane potential. For testing high and low ion uptake performance, defined concentrations of N, P, K and S were added to the basal solution (see Materials and Methods). Plants were then transferred to individual hydroponic chambers containing the nutrient solution, and the solution was sampled over time using a coupled 24-channel peristaltic pump. The net nutrient uptake rate was determined from the depletion of the nutrient from the chamber over time. The nutrient solution samples were collected in a collection plate and the ion concentrations were quantified using ion chromatography (Fig. 1*F*). The downstream data analysis for calculating specific nutrient uptake rates was automated using R scripts (Fig. 1*G* and *H*) (https://doi.org/10.5281/zenodo.3893945).

The multiple ion uptake pipeline was used to determine 35 uptake parameters for each plant (described in Table S1). Including all plant traits measured, a total of 50 traits were collected for each plant. These traits were used to determine the nutrient uptake capabilities of each plant in terms of: (i) the total uptake performance of a plant, (ii) the specific ion uptake rate on a root length or mass basis, and (iii) the ion uptake ratio and stoichiometry of the plant. The pipeline is modular and flexible for adoption with different plant species by changing vessel volumes, nutrient concentrations, and experiment designs. The phenotyping platform enables exploration of a broad range of questions, including studies of competitive and facilitative interactions between nutrients, influence of abiotic and biotic factors on nutrient uptake, and the use of mutants for genetic confirmation of nutrient uptake properties.

### Genetic diversity exists among NAM population founder lines for specific nutrient uptake rates and is highly heritable

Using the multiple ion uptake phenotyping pipeline, specific nutrient uptake rates of nitrate, ammonium, phosphate, potassium, and sulfate were characterized simultaneously for the 25 maize NAM population founder lines and the reference line B73. After growth in a common nutrient solution, each line was deprived of the focal macronutrients for 48 hours, and then transferred to the uptake setup. Nutrient uptake performance was characterized in a high and a low multiple ion solution with a 10-fold macronutrient concentration difference to characterize high and low affinity performance (23, 33, 34).

We found that genetic diversity exists among the NAM population founders for specific nutrient uptake rates (on a root length basis), with significant genotypic differences for nitrate (P < 0.001, Fig. 2*A* and *B*, Table S2) as well as for phosphate, potassium, sulfate and ammonium (P < 0.01, Fig. S1, Table S2). Line M162W had the greatest specific nitrate uptake rate in the high concentration solution and second highest in the low concentration solution, with 4.81 and 2.37 times that of the lowest line CML69, respectively (Fig. 2*A* and *B*). Furthermore, significant effects of genotype × concentration were detected for nitrate and sulfate (P < 0.05, Table S2). The founders showed differential nitrate uptake rate capacity between the concentration solutions, with the greatest difference observed for line Ki3, which exhibited a specific uptake rate in the high concentration that was 5.29 times higher than in the low concentration. Line HP301 appeared to have a near maximum specific uptake rate (I_max_) in the low concentration as its ratio of uptake between the two nutrient concentrations was 1.1, highlighting the broad environmental acclimation of the population (Fig. 2*C*).

**Fig. 2.**
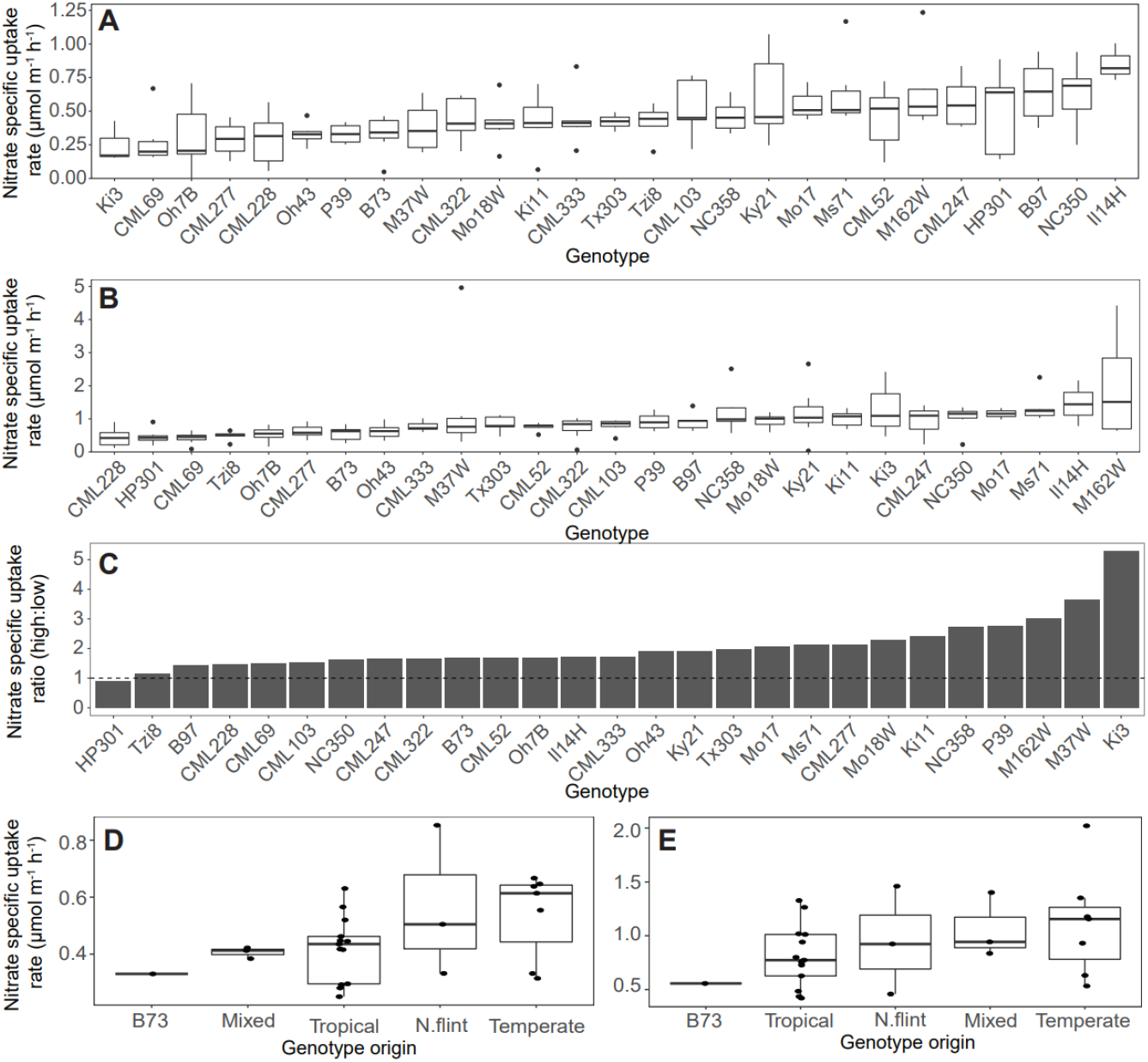
Genetic diversity among NAM population founder lines for specific nitrate uptake rates from solution concentrations of (A) 100 μM and (B) 1 mM (P < 0.001). (C) The net specific nitrate uptake rate ratio between the solutions, with a ratio greater than 1 representing a greater uptake rate in the high concentration compared to the low concentration. The specific nitrate uptake rates by broad group classification in the solution concentrations of (D) 100 μM and (E) 1 mM. Figures for phosphate, potassium, sulfate and ammonium are available in Fig. S1.

The specific nutrient uptake rates for each macronutrient were found to be highly heritable with a broad-sense heritability between 0.37 and 0.88 (Fig. S2). Therefore, the traits assessed by this pipeline may serve as novel breeding targets with genetic variation that could be harnessed for breeding crops better able to acquire soil nutrients. Based on broad group classification of the NAM population founders by phenology and breeding background, the specific nutrient uptake rates were compared (Fig. 3*D* and *E*) (35). Uptake performance in both high and low concentration solutions were found to not be significantly different among broad classification groups (P = ns) and, therefore, specific nutrient uptake rate has greater variation by line rather than origin.

**Fig. 3.**
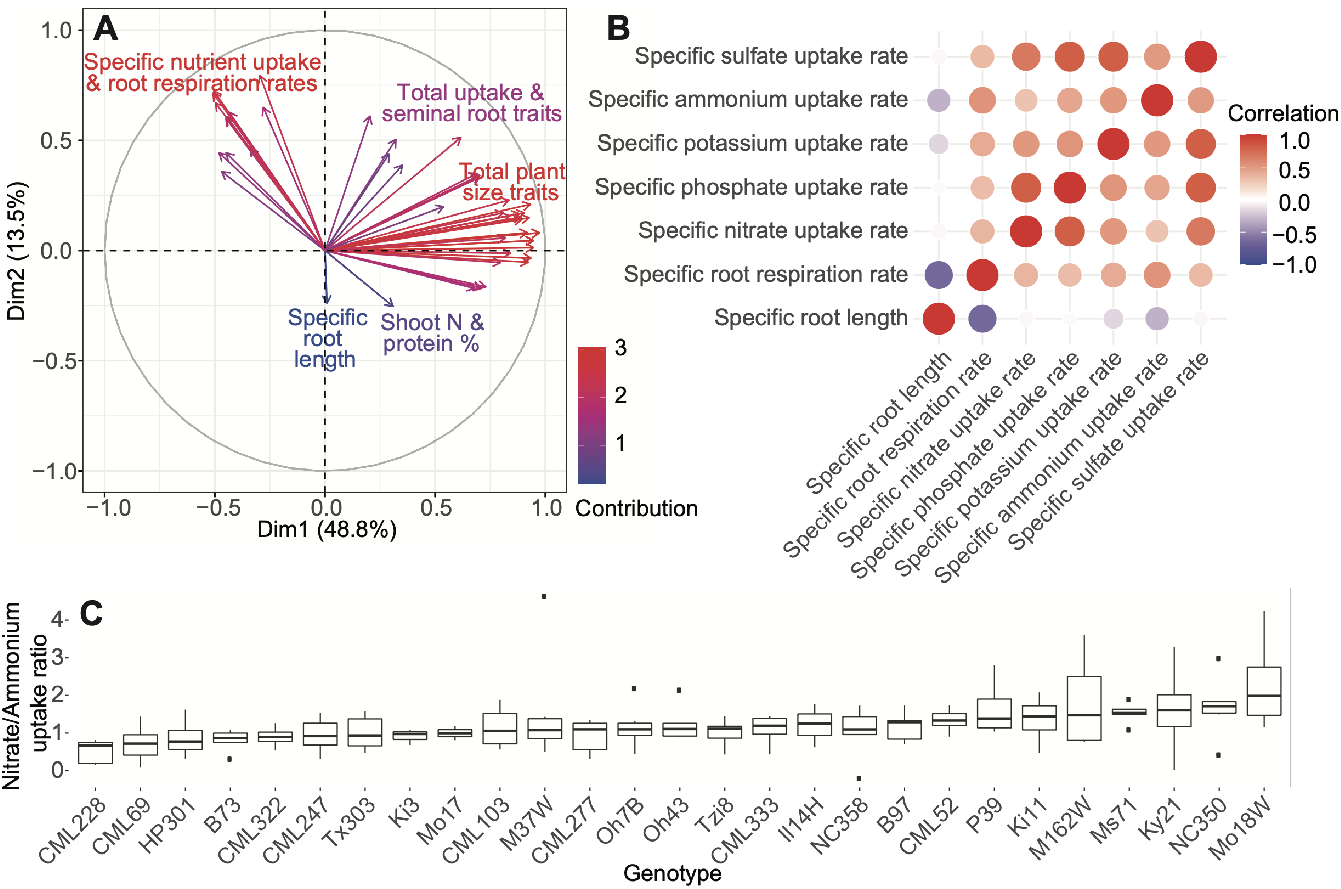
(A) PCA ordination of extracted plant traits of the NAM population founder lines in the high concentration solution. Arrows indicate directions of loadings for each trait and are color coded by contribution to the percent variation of the component. (B) Correlation matrix for specific root nutrient uptake, respiration and length parameters in the high concentration solution. Correlations are color coded from strong positive correlation in red to strong negative correlation in blue with no correlation shown in white. (C) The net uptake rate ratio between nitrate and ammonium in the 1 mM high concentration solution (P < 0.001). A ratio above 1 represents a higher proportion of ammonium uptake compared to nitrate. ANOVA results for all nutrient combinations are shown in Table S2.

To determine if specific nutrient uptake rate in a plant is relative to a particular nutrient concentration, or consistent across a wide concentration range, a regression between the specific nutrient uptake rates by concentration was made. A positive significant relationship was observed between the concentration levels for specific nutrient uptake rates of nitrate (P < 0.05, Fig. S3), with a non-significant trend for sulfate (P = 0.08, Fig. S3). For phosphorus, potassium and ammonium, however, no significant relationship was found between specific uptake rates in the two concentrations (P = ns, Fig. S3). Therefore, measurements in a single nutrient concentration representative of dominant field conditions at the respective growth stage may have predictive power for crop selection. Wide genotypic variation in specific nutrient uptake rate was observed within each nutrient, representing value for screening lines at multiple nutrient concentrations for more detailed characterization when possible.

### Specific nutrient uptake rates among multiple macronutrients are correlated and are a distinct efficiency breeding target

The overall interactions among plant traits were determined using the genetically diverse NAM population founder dataset. A principal component analysis (PCA) for all 50 uptake and plant was conducted, which showed that the first two PCs accounted for more than half (62.3%) of the total variance. The PCA ordination revealed a distinct separation between the specific nutrient uptake and root respiration rates from the total uptake and plant size measures (Fig. 3*A*). This trait separation indicates independent genetic control and, therefore, distinct breeding targets. As specific nutrient uptake and respiration rate traits are independent of plant size, the ratio of these two traits could be considered as an index of efficiency.

A correlation matrix of the specific nutrient uptake rate data revealed positive correlations among nutrient types (correlation range 0.29 – 0.80, P < 0.001, Fig. 3*B*). In the high concentration, the correlations between nitrate:phosphate, sulfate:phosphate, and sulfate:potassium were the highest with each having correlation scores of 0.80. The correlation scores between the specific nutrient uptake rates in the low concentrations were generally lower than in the high concentrations (0.11 – 0.69, P < 0.01) and with no significant correlations between nitrate:ammonium, potassium:ammonium, and sulfate:ammonium (P = ns). The exceptions were nitrate:potassium and ammonium:phosphate, which had higher correlations in the low than the high concentration (0.68 and 0.51, respectively, P < 0.001, Fig. S4). The positive correlations among specific macronutrient uptake rates represent shared underlying mechanisms of regulation for uptake.

Interestingly, despite the overall trend of positive correlation among specific nutrient uptake rates, we observed that there is also genotypic variation in nutrient stoichiometry. Uptake ratios between nutrients were unequal amongst the lines, with preferences exhibited for particular nutrient combinations (Fig. 3*C*). Significant genotypic differences in specific nutrient uptake rate ratios were observed between the nitrogen forms nitrate and ammonium (P < 0.01, (Fig. 3*C*), and phosphate and potassium (P < 0.01, Fig. S5). The line with the greatest preference for ammonium over nitrate was Ms71, having 2.61 times higher specific uptake rate of ammonium compared to nitrate. In contrast, CML228 had a nitrate preference over ammonium with a specific nutrient uptake rate ratio of 0.03. Therefore, genotype selection tailored to a particular soil nutrient composition and management strategy has the potential to improve uptake efficiency and yield.

### Maize lines with greater low affinity specific nitrate uptake have higher specific root respiration to facilitate active transport

To examine the metabolic costs associated with nutrient uptake, we measured the specific root respiration rates for each of the NAM population founder lines immediately after the uptake experiment. The specific nitrate, potassium and sulfate uptake rates were found to have a significant positive correlation with specific root respiration in the high concentration solution only (R = 0.12, 0.16, 0.15, P < 0.001, Fig. 4*A* and *B*), Fig. S6). That the correlation between specific nutrient uptake rate and respiration rate was only observed in the high concentration indicates that high root respiration itself is not causative of increased uptake (P = ns, Fig. 4*A* and *B*). When exposed to the high concentration an average increase of 27% in specific root respiration rate was observed across all lines. Lines with a greater uptake capacity likely respired more to facilitate the active transport of nutrients for a higher specific uptake rate (Fig. 4*B*). For phosphate and ammonium, positive correlations between specific root respiration rates and specific nutrient uptake rates were observed in both nutrient concentration solutions (R = 0.16, 0.24, P < 0.001, Fig. S6). These results indicate that increased respiration may be a cost of increased uptake, but that simultaneously selecting for higher uptake but lower respiration may be possible.

**Fig. 4.**
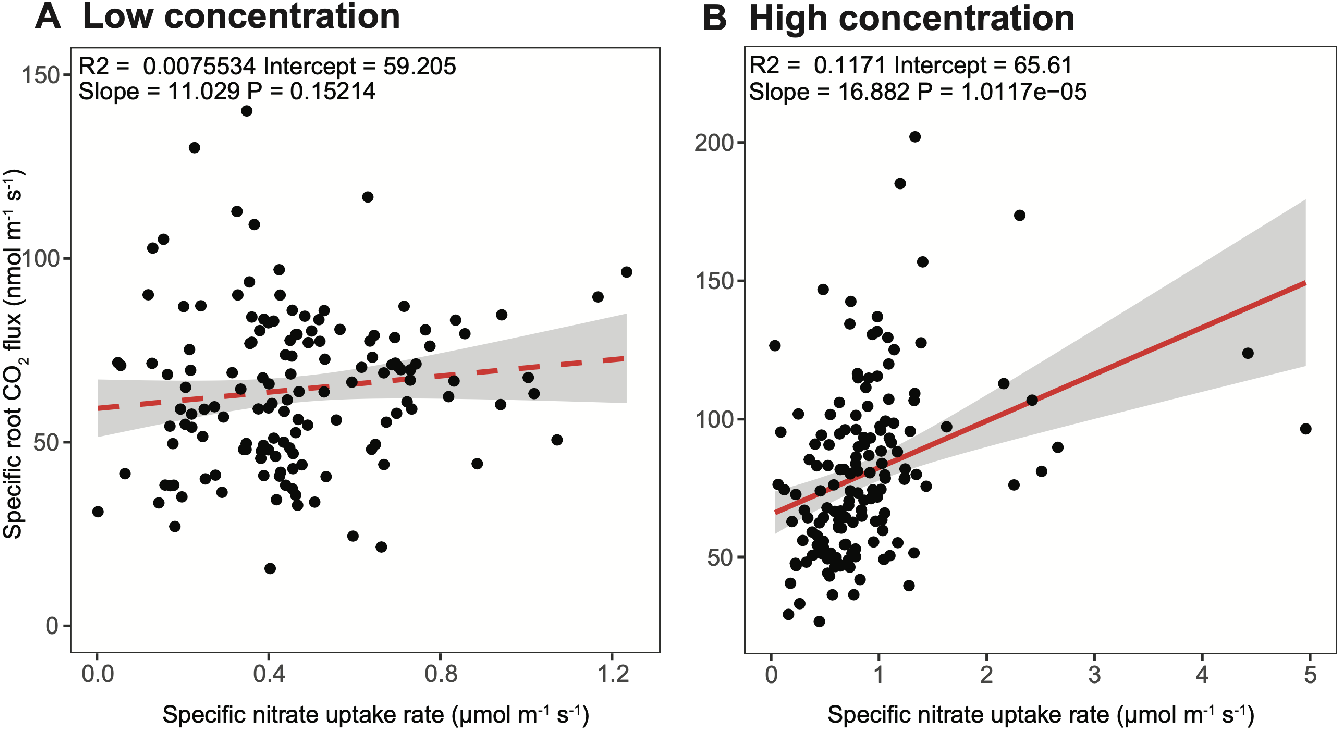
Linear regression analyses for the NAM population founder lines between specific nitrate uptake rate and specific root respiration rate in the (A) low (100 *μ*M) and (B) high (1 mM) concentration solutions. Significant relationships are depicted with a full red line and non-significant relationships with a dashed red line. The grey bar represents a 95% confidence region.

### Lines with high specific nutrient uptake rate have elevated transcript abundance of nutrient responsive genes and metabolism

To elucidate the pathways and differences between lines with high and low specific macronutrient uptake rates at the molecular level, a comparative transcriptomic analysis was conducted. Two lines with high specific uptake rates (M162W and Ky21), three lines with low specific uptake rates (CML69, CML227 and CML228) and the reference line B73, which showed an intermediate specific uptake rate in both low and high concentration solutions, were selected for RNA-seq (Fig. 5*A* and *B*). The phenotyping experiment described above and most experiments in the literature include a deprivation step generally believed to increase uptake rates by priming the molecular machinery. To understand short-term transcriptomic responses specific to the high nutrient uptake rate lines, maize seedlings were grown in full nutrient conditions for 12 days and then one half of the seedlings were macronutrient deprived for 48 hours, which was the same procedure used for the phenotyping experiment. We hypothesized that both NAM lines with high specific nutrient uptake rates (Ky21 and M162W) under nutrient deprived conditions utilized a conserved set of pathways and genes to mediate elevated nutrient uptake that were reduced or missing in NAM lines which were unable to do so (CML69, CML227, CML228 and B73).

**Fig. 5.**
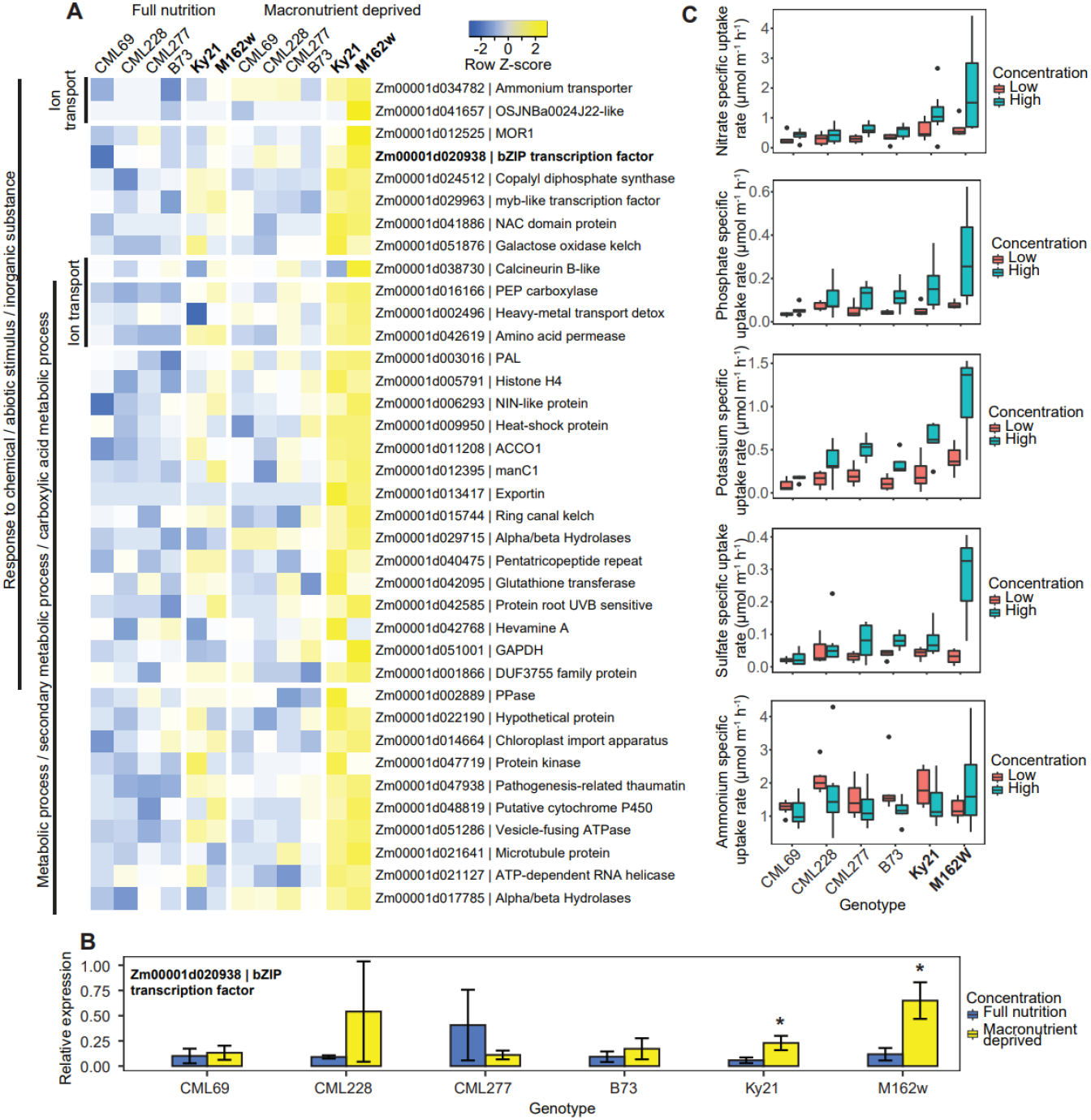
Maize founder lines showing differences in root gene expression across full nutrient and deprived plants. (A) Heatmap showing selected genes induced during full nutrient and nutrient deprived conditions with fold-changes of ± 1 (P < 0.06). Z-score represents the number of standard deviations of a condition from the mean of all conditions. (B) qPCR validation results of candidate bZIP transcription factor across full nutrient and deprived plants (P < 0.05). (C) Specific nutrient uptake rates of selected lines used in RNA-seq by nutrient solution concentration.

In our transcriptional profiling, the dataset was analyzed to specifically identify responses in common among high specific nutrient uptake rate lines but absent across all other lines under both full and deprived conditions. A subset of 50 genes that were significantly upregulated and 182 genes that were downregulated by at least ± 1 log fold-change (P < 0.06) were selected by comparing the gene expression of high specific nutrient uptake lines in low nutrient conditions with the expression of genes in all other samples (Fig. 5*A*, Data S1). Gene Ontology (GO) enrichment analyses revealed a significant activity of metabolism related genes (GO:0044710, 64%), corroborating the higher root respiration rates seen in these lines (Fig. 5*A*, Data S2). A second major GO enrichment category was response to chemical / abiotic stimulus / inorganic substance (GO:0042221, 47% / GO:0009628, 34% / GO:0010035, 26%), supporting the enhanced nutrient deprivation response in the high specific uptake rate lines.

Compared to B73 and the low specific nutrient uptake rate lines, genes encoding a number of nutrient transporters were found in the high specific nutrient uptake rate lines (Fig. 5*A*, Data S1). These included an ammonium transporter (Zm00001d034782) and the NIN-like protein transcription factor (Zm00001d006293), supporting our observations of higher uptake of these nutrients in these lines. This transcription factor was recently shown to be a central regulator of nutrient-signalling networks (36). These findings validate our discovery platform and analyses by selecting candidate lines based on specific nutrient uptake rate performance. Our data suggest that this transcription factor traditionally thought to mediate nitrate uptake and metabolism might be involved in fundamental multiple-nutrient uptake or metabolism processes. Further functional studies are warranted to confirm this hypothesis.

Finally, we found a number of novel targets that, with further investigation, may turn out to be useful to improve nutrient uptake performance. One of the novel genes validated with qPCR was Zm00001d020938, a bZIP transcription factor family protein that was induced during nutrient deprivation in the high nutrient uptake rate lines only (P < 0.05, Fig. 5*B*). The transcription factor is likely a regulator of downstream signalling cascades mediating higher nutrient uptake or metabolism (Fig. 5*A*, Data S2). These gene candidates could potentially provide breeding targets for maize lines with greater nutrient uptake efficiency.

## Discussion

With the wide adoption of image-based root phenotyping in recent years, significant advances in characterizing root system architecture have been made (37, 38). However, understanding of functional root processes including nutrient uptake lags behind, with significant knowledge gaps remaining about the genetic, physical, and molecular mechanisms involved. Development and adoption of phenotyping approaches for uptake kinetics scaled to mapping populations could accelerate this discovery (reviewed by (25)).

Our approach addresses this challenging bottleneck with the development of a modular pipeline for reproducible high-throughput phenotyping of multiple ion uptake by roots. In maize, we revealed that specific nutrient uptake rate (nutrient uptake rate per length of root) is an uptake efficiency trait in respect to root construction costs that is distinct from total plant uptake and plant size traits (Fig. 6*A* and *B*). Specific ion uptake rates for several macronutrients were found to be highly heritable and variable among the genetically diverse NAM population founder lines. Harnessing this natural variation through identification of underlying genes and mechanisms is of paramount importance to improving nutrient uptake efficiency in crops. Work in maize indicated variation in uptake kinetics even among root classes such as seminal, nodal, and lateral roots, which implies that regulation of transporters and other molecular machinery leads to substantial differences in uptake (39). Allelic variation of a nitrate transporter affecting specific uptake rate in rice demonstrates that a single allele can significantly affect plant resource acquisition (40). Recently, mining natural sequences for more effective RuBisCO alleles led to discovery of variants with six-fold faster reactions than typical plant variants (41), and similar strategies could be used for nutrient uptake. Breeding efforts for yield may have indirectly selected for increased specific nitrogen uptake rate in modern wheat varieties (42) and, therefore, crop selection for specific ion uptake rate directly could possibly accelerate gains in nutrient uptake efficiency and yield.

**Fig. 6.**
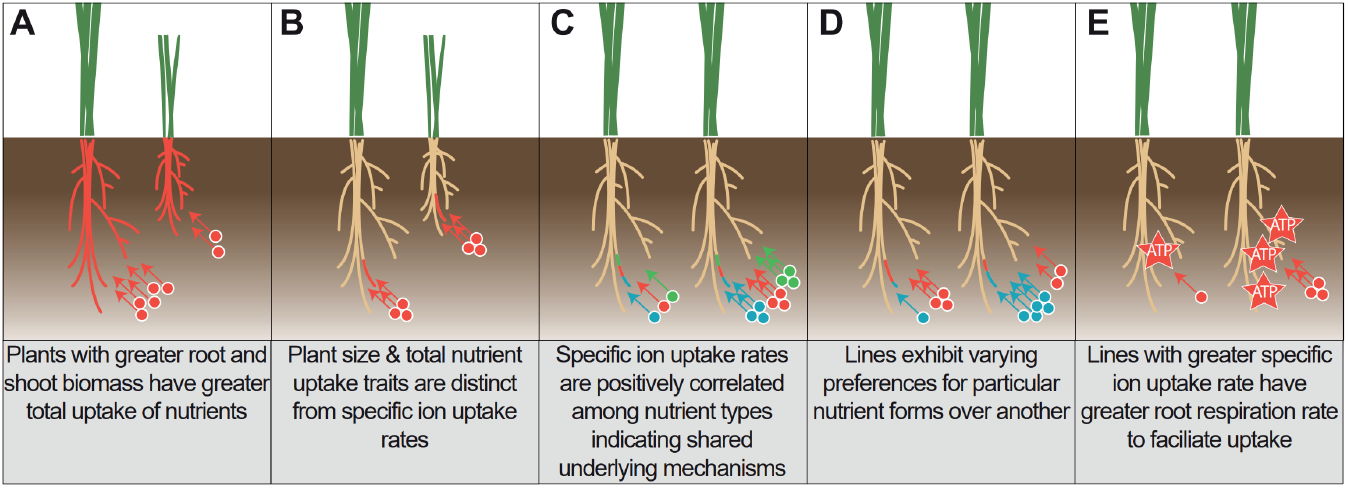
Mechanisms of uptake performance with regards to plant size, specific uptake rates, uptake of different nutrients, and respiration. Root colors represent the area of root nutrient uptake per nutrient type. Circle colors represent different nutrient types.

The multiple ion phenotyping approach allowed investigation of the interaction between nutrients in plant uptake. We uncovered that specific ion uptake rates are positively correlated among the macronutrients N, P, K, and S and, therefore, are likely governed by shared mechanisms (Fig. 6*C*). Only a few studies have measured the uptake rates of more than one nutrient, and even more rarely investigated in the context of discovering shared mechanisms through correlative analysis across lines (26, 32). Nutrient transporters have been shown to exhibit cross-regulation with multiple nutrients at the local and whole plant levels as well as to facilitate uptake of phytohormones (43, 44). Our RNA-seq analyses identified central regulators of nutrient-signaling networks that have elevated transcript abundance in the high specific nutrient uptake lines.

With the current push towards multi-dimensional phenomics (45–47), conducting nutrient uptake experiments in representative conditions is important to assess the complexity of interplay between nutrients. Adding to this complexity, we also found significant variation in preference for specific nutrients in particular nutrient combinations among the NAM population founder lines (Fig. 6*D*). This illustrates the importance of characterizing cultivars to ensure that they are adapted to the soil environment and fertilizer regimes according to nutrient uptake characteristics. Dissection of these mechanisms is important to understand the fundamental processes by which all plants forage nutrients from their environment. This study used a hydroponics approach as it has inherent practical advantages over soil in controlling nutrient availability and measurements of uptake rates. It will be important to extend this research to determine how soil physical and chemical properties affect multiple nutrient uptake rates and nutrient preferences. Optimization of above- and below-ground plant traits to the environment and management practices will be integral to improving nutrient uptake and reducing fertilizer losses.

Nutrient uptake is a substantial carbon expense to the plant and, therefore, understanding how nutrient uptake affects overall plant efficiencies is vital. We phenotyped the specific root respiration rates amongst the NAM population founder lines as a measure of root activity and metabolic cost. The results revealed that specific nutrient uptake rates were positively correlated with specific root respiration rate in the high nutrient concentration solutions (Fig. 6*E*). As specific root respiration rate was not correlated in the low concentration solution for nitrate, potassium and sulfate, the metabolic cost appears to be associated with low affinity transport of these nutrients. Potentially, respiration is linked to uptake capacity by co-regulation of associated genes and the need of ATPase and other pumps to form necessary ionic gradients. The large variation observed in the NAM population founder lines for specific nutrient uptake rate likely represents the diverse environments to which they are adapted. Our RNA-seq data revealed a significant enrichment of metabolism-related genes in the high specific nutrient uptake lines, which may corroborate the enhanced nutrient uptake rates observed. The positive relationship between uptake and respiration raises an interesting dilemma. On the one hand, maximizing uptake should generally be beneficial (25), while on the other, minimizing metabolic burden of the root system has been proposed to be a promising opportunity (48). The *SimRoot* simulation model generally indicated that increasing uptake rates was beneficial, but since no cost is included in the model the optimum uptake rate may actually be slower (39). Including accurate costs in such simulations will greatly facilitate efforts to design ideal integrated phenotypes. One way to attain reduced root respiration is with a greater percent of root cortical senescence, although this was found to also lower N and P uptake (49) and other anatomical traits may have similar influences. Therefore, we propose that measuring both uptake and respiration rates is necessary in order to ensure co-optimization.

Our results highlight the importance of nutrient interplay and that representative nutrient uptake assays should be in the presence of other nutrients as is the case in soil. The multiple ion uptake platform, *‘RhizoFlux’*, enables high-throughput and precise phenotyping, which will provide mechanistic insights into nutrient uptake and has a great potential for genomic selection that will benefit agriculture. The results revealed that specific ion uptake rates are highly heritable and, therefore, we envision that breeding for targeted environments by combining above- and below-ground plant traits to form integrated phenotypes will likely improve plant performance and yield whilst reducing fertilizer losses.

## Materials and Methods

Material dimensions are given in the units supplied by the manufacturer.

### Plant materials

Seed for the maize Nested Association Mapping (NAM) population founder lines and reference line B73 were obtained from Dr. Felix Fritschi (University of Missouri, originally sourced from Dr. Sherry Flint-Garcia, USDA-ARS). The founder lines were originally selected to maximize diversity from a larger panel of diverse maize inbreds, and each has a recombinant inbred population crossed with the common reference parent B73.

### Experimental design and growth conditions

The experiment was a complete randomised block design replicated seven times over time as independent runs. Two lines had poor germination that reduced their sample number, Mo17 (two replicates high and three replicates low) and Ki3 (three replicates each). Seeds were surface sterilized with 5% bleach and washed three times with double deionized water (ddH_2_O). Sterilized seeds were transferred to germination paper rolls soaked with 0.2 mM CaSO_4_, and then allowed to germinate at 28 °C for four days in the dark (Fig. 1*A*). Uniformly germinated seedlings were transferred to individual plastic mesh plant baskets (1.5” × 2”, Shenzhen Skywalker Electronic Limited, Shenzhen, China) that were placed into plant cone-tainers (SC10 Super RL98 cell, Stuewe Sons Inc., OR, USA). A slot about 3 mm wide was cut from the bottom of the cone to about 5 cm from the top in order to accommodate the tubing from the sampling platform described below. The plants in cones were placed in aerated hydroponic tanks fitted with custom acrylic lids with 24 equally spaced holes of 1.75” diameter such that the cones were held vertically (Dividable Grid Container 10.88” × 16.5” × 8”, Quantum Storage Systems, FL, USA; EcoPlus commercial air pump, Hawthorne Gardening Company, WA, USA) (Fig. 1*B-D*). The nutrient solution used was a modified ½-strength Hoagland’s solution composed of (in *μ*M) 500KH_2_PO, 5700 KNO_3_, 300NH_4_NO_3_, 2000CaCl_2_, 1000MgSO_4_, 46H_3_BO_3_, 7ZnSO_4_ · _7_H_2_O, 9 MnCl_2_ · _4_H_2_O, 0.32 CuSO_4_ · _5_H_2_O, 0.114 (NH_4_)_6_Mo_7_O_24_ · _4_H_2_O, and 150Fe(III) EDTA(C_10_H_12_N_2_NaFeO_8_). Additional Fe(III)-EDTA was added every three days and the solution was adjusted to pH6 using chHCl. Plants were grown in a growth chamber with a day:night cycle of 16/8 h at 28/20°C at a photon flux density of 400 *μ*mol^-2^ s^-1^ at canopy height for 12 days (E7/2 growth chamber, Conviron, Winnipeg, Canada). The plants were then transferred to a largely macronutrient-free nutrient solution composed of (in *μ*M, (500CaCl_2_, 46H_3_BO_3_, 7ZnSO_4_ · _7_H_2_O, 9 MnCl_2_ · _4_H_2_O, 0.32 CuSO_4_ · _5_H_2_O, 0.114 (NH_4_)_6_Mo_7_O_24_ · _4_H_2_O, and 150Fe(III) EDTA(C_10_H_12_N_2_NaFeO_8_) for 48 h before measurement of multiple ion uptake.

### Multiple ion uptake and trait measures

For the ion uptake phenotyping assay a modular platform was developed with individual plant hydroponic chamber control of nutrient solutions (Fig. 1*E*). A single module consisted of 24 polyvinyl chloride (PVC) pipe chambers (1.5” ID PVC Schedule 40 pipe and 1.5” hub cap fitting, Charlotte Pipe and Foundry Company, NC, USA) with a volume capacity of 250 mL. The chambers were designed so each plant could remain in the seedling cone for minimal disturbance to the roots during transfer. Each chamber was connected to two 24-channel peristaltic pumps (Ismatec ISM944A, Cole-Parmer Instrument Company LLC., IL, USA) with tubing. One pump was used to fill the empty chambers with nutrient solution and afterwards provided continuous aeration by pumping air to the chambers (24 rotations per minute) via tubing connected to the bottom of each chamber (3.17 mm ID tubing Ismatec SC0222-LT 2-Stop 0; Masterflex SC0223-LT Tygon; Masterflex Hose Barb Union 1/8”, Cole-Parmer Instrument Company LLC., IL, USA; CPC ID 1/8” hose barb PMCD1702 insert PMC2202, Colder Products Company, MN, USA). To fill with nutrient solution, the 24 tube inlets were placed into a common container full of the appropriate solution, the pump ran for a predetermined time to output the correct solution volume, and then the tubes were removed to pump air for aeration. The second pump was used for periodic sampling of the nutrient solution from the middle of the chamber into a 2 mL 96-well collection plate (0.51 mm ID tubing Ismatec SC0005-LT 2-Stop 0; Masterflex SC0029-LT Tygon, Cole-Parmer Instrument Company LLC., IL, USA; Diba MicroBarb^®^ Adapter, 1/4” to 0.02” ID Diba Industries Inc., CT, USA). The correct sample volume was achieved by running the pump for a predetermined time. After each nutrient sampling, the pumping direction was reversed to expel all solution back into the chamber and clear the tubing. The small diameter tube outlets were placed into a 96-well microplate cover (VWR International, LLC, PA, USA) with holes drilled in a pattern to match the 96-well collection plate. Two identical modules with a total of four pumps were used and placed in a large growth chamber with the same environmental conditions as detailed earlier (PGR15 growth chamber, Conviron, Winnipeg, Canada) with a throughput of 48 plants per experimental run.

For the NAM population founder lines, the ion uptake assay was used to phenotype multiple ion uptake kinetics under high and low macronutrient concentration solutions. The high concentration solution consisted of (in *μ*M) 1000KNO_3_, 1000NH_4_Cl, 125Ca(H_2_PO_4_)2H_2_θ, 250MgSO_4_, and 375CaCl_2_. For the low concentration solution, the macronutrient concentrations were 10-fold lower than the high solution with (in *μ*M) 100KNO_3_, 100NH_4_Cl, 12.5Ca(H_2_PO_4_)_2_H_2_O, 25MgSO_4_ and 487.5CaCl_2_. In both solutions, the calcium concentration was maintained at 0.5 mM and a 1 mM MES buffer was used (pH 6). As noted above, plants were grown for 12 days in complete nutrient solution at relatively high concentrations, then underwent deprivation in a nutrient solution lacking macronutrients for 48 h. Based on preliminary work during development, a nutrient induction step was not used as no significant effect on specific nutrient uptake rate was observed across nutrient concentration ranges (10 *μ*M-10 mM). The plants were transferred intact within their plastic cones into individual chambers in the phenotyping platform such that the slot in the cone fit over the inlet of the sampling tube. Once the first pump filled all 24 chambers with the appropriate solution, all plants were transferred to the chambers with the plastic cones lowered into the chambers and submerging the roots. Two minutes after the macronutrient deprived plants were transferred to the individual chambers the first 1.5 mL nutrient sample was collected for time zero. Nutrient solution samples were then taken at 0.5, 1, 2, 3, 4, 6 and 8 h.

The ion concentrations of the collected nutrient samples were determined using a Thermo Scientific ICS-5000+ ion chromatographic system (Thermo Fisher Scientific, MA, USA). Chromatographic separation was achieved using a Dionex IonPac CS12A (2 × 250 mm) analytical column with a AG12A (2 × 50 mm) guard column for cations, and a Dionex IonPac AS11HC-4 m (2 × 250 mm) analytical column with a AG11HC-4 m (2 × 50 mm) guard column for anions. Ions were eluted using gradient elution at a flow rate of 0.3 mL min^-1^ for cations and 0.33 mL min^-1^ for anions and detected by a self-regenerating suppressor and a conductivity detector. Column temperature was maintained at 20.5 °C and the injection volume was 25 *μ*L. The cation eluent source was a Thermo Scientific Dionex EGC III Methanesulfonic acid eluent generator cartridge. Elution of cations was achieved with the following gradient: 12 mM to 20 mM in 7 minutes, held at 20 mM for 8 minutes, ramped from 20 mM to 40 mM in 3 minutes, the column was re-equilibrated at 12 mM for 5 minutes. The anion eluent source was a Thermo Scientific Dionex EGC KOH cartridge. Elution of anions was achieved with the following concentration gradient: 6 mM to 21.5 mM in 16.5 minutes, 21.5 to 60 mM in 6.5 minutes and held at 60 mM for 3 minutes, the column was re-equilibrated at 6 mM for 8 minutes. Standards for the cations (Thermo Scientific Dionex Six Cation-II) and anions (Thermo Scientific Dionex Seven Anion Standard II) were used and the data were extracted using the Chromeleon 7.2 SR4 software (Thermo Fisher Scientific, MA, USA).

Immediately after the final uptake assay sample collection, the roots were severed from the shoots and root respiration for each plant was measured. Roots were transferred into a 43 mL airtight chamber connected to the LI-8100 Automated Soil CO_2_ Flux System (LI-COR Biosciences, NE, USA). The CO_2_ flux in the chamber was then measured with an observation duration of 90 seconds and dead band set at 20 seconds using the LI-8100A v4.0.9 software. Total respiration rate was calculated automatically by the linear fit mode in SoilFluxPro v4.0.1 software with a curve fit time of 20-90 seconds and 0.1 soil area. After root respiration, the root system was stored at 4 °C in 70% ethanol for later imaging using a flatbed scanner equipped with a transparency unit (Epson Expression 12000XL, Epson America Inc, CA, USA). Roots were spread out on a transparent Plexiglas tray with a 5-mm layer of water and imaged at a resolution of 600 dpi. The seminal, lateral and secondary-order lateral root lengths for each plant were calculated from the flatbed images using WinRhizo™software 2013e (Regent Instruments Inc., Canada) based on diameter thresholds (in mm) of 0.8-4.25, 0.15-0.8 and 0-0.15, respectively. The leaves were separated from the stems and laid out on a custom leaf vice made from two sheets of Perspex, and then imaged at a resolution of 300 dpi using a flatbed scanner equipped with a transparency unit. The leaf length and area for each plant was determined from the images using a custom imageJ macro (https://doi.org/10.5281/zenodo.3893945) modified from (50). The root system, leaves and stems were then dried at 60 °C for 3 days for determination of dry weights.

For determination of root and leaf N contents, the root and leaf dry matter were ground into powder by placing the samples into glass vials with three opposing blades and shaking at a frequency of 30 Hz for 10 minutes using a Qiagen TissueLyser II (Qiagen, ML, USA). Ground root and leaf percent N was determined by the Dumas method using the Elementar rapid N exceed analyzer (Elementar Americas Inc, NY, USA). Samples were weighed into tin foil sample papers (Elementar Americas Inc, NY, USA) without any pre-treatment. Samples were run using a standard method implemented in the instrument software, with a total analysis time of about 5 minutes. CO_2_ was used as the carrier gas and L-aspartic acid (Sigma-Aldrich, St. Louis, MO, USA) was used as a standard. A nitrogen-to-protein content conversion factor of 6.25 was applied to calculate the average protein content (51). Rapid N Exceed software V.1.1.26 (Elementar Americas Inc, NY, USA) was used for data processing.

### Transcriptomic Analysis

The entire root system was collected from maize seedlings grown in hydroponics. Seedlings were grown in full nutrient conditions for 12 days and then one half of the seedlings were macronutrient deprived for 48 h whilst the other half remained in full nutrient solution. The macronutrient deprivation was the same as used in the phenotyping experiment. Three biological replicates each consisting of 2-3 plants per line per treatment were collected. Samples were immediately frozen at 70 °C and later ground with a pestle and mortar under liquid nitrogen. Total RNA was extracted from the frozen tissues using Spectrum™Plant Total RNA Kit (Sigma-Aldrich, St. Louis, MO, USA) following the manufacturer’s instructions. RNA quality was checked with Agilent Bioanalyzer 2100 (Agilent, Palo Alto, CA, USA) and quantified using Qubit™RNA BR Assay Kit (Thermo Fisher Scientific, MA, USA). One *μ*g of DNase-treated total RNA was used for library construction using TruSeq Stranded mRNA Library Prep Kit (Illumina Inc, CA, USA) following the manufacturer’s protocol. Library quality was checked using TapeStation (Agilent, Palo Alto, CA, USA) and quantified by Qubit™RNA BR Assay Kit (Thermo Fisher Scientific, MA, USA). Each library was sequenced at 150 bp paired end at 30-40 million reads using an Illumina Hiseq sequencer (Novogene Co Ltd, Beijing, China).

The RNA-seq data set was mapped against the maize reference genome B73 RefGen V4 AGPv4 (https://maizegdb.org/genome/assembly/Zm-B73-REFERENCE-GRAMENE-4.0) (52). Gene expression was quantified as Fragments Per Kilobase Of Exon Per Million Fragments Mapped (FPKM), and differentially expressed genes with a false discovery rate of less than 0.1 were accepted (53). Differential gene expression analysis was performed using DESeq2 with DEBrowser V1.17.1 by comparing gene expression of high specific nutrient uptake lines in low nutrient conditions to expression of genes in all other samples (53, 54). Genes with a maximum count of less than 10 across all samples were filtered out and the data was normalized using Median Ratio Normalization (MRN). Gene log fold-change was rounded to two decimal places and candidate genes were selected based on a ± 1 log fold-change criterion with a P value less than 0.06. Gene ontology (GO) enrichment analysis was performed with maize reference genome B73 RefGen V4 AGPv4 using Agrigo V2 (55). The expression changes of candidate genes were plotted using Expression Heatmapper tool (56).

Primers used to amplify candidate macronutrient deprivation-responsive gene transcripts were designed by determining the exon regions for each gene using Gene Structure Display Server 2.0 (40) and logged using Geneious software (Biomatters Ltd, Auckland, NZ). Primers were designed using the last exon of each gene, avoiding primers with predicted hairpins where possible, using Primer3 v4.0.0 software (57). Primer-BLAST was used to confirm the specificity of primer pairs for the intended targets. Clustal omega (58) was used to confirm that primers should bind to all splice variants of gene transcripts. For subsequent reverse transcription-quantitative PCR (RT-qPCR), 5 *μ*g of total RNA was treated with TURBO DNA-free™Kit (Invitrogen, Thermo Fisher Scientific, MA, USA) to remove any potential genomic DNA contaminants. Two *μ*g of DNA-free total RNA was used for first-strand cDNA synthesis using SuperScript™III reverse transcriptase (Invitrogen, Thermo Fisher Scientific, MA, USA). qRT-PCR was performed with KiCqStart™SYBR^®^Green qPCR ReadyMix™(Sigma-Aldrich, St. Louis, MO, USA) using QuantStudio™ 7 Flex Real-Time PCR System (Applied Biosystems, Thermo Fisher Scientific, MA, USA). The primer pairs used are listed in Table S3. Data were collected and analyzed using the QuantStudio™7 Flex Software (Applied Biosystems, Thermo Fisher Scientific, MA, USA). Differential gene expression was quantified based on the △ △Ct method using normalized geo-metric means of the two reference genes (Zm00001d002944, Zm00001d020826; (59)).

### Statistical analysis

Statistical analyses were conducted using R version 3.6.0 (60); the statistical analysis R codes including the packages needed are available (https://doi.org/10.5281/zenodo.3893945).

The depletion rate of a nutrient from a solution is commonly accepted as equal to the net uptake rate by roots (assuming both influx and efflux). Therefore, the following equation was used to determine the total net influx rates for nitrate, ammonium, potassium, phosphate and sulfate:

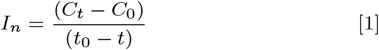

where *I_n_* is the net influx into the plant; *C*_0_ is the initial concentration of the solution at the start of the experiment *t*_0_; *C_t_* is the concentration at sampling time. The In was then divided by either the root system length (cm) or weight (g) to calculate the net specific nutrient uptake rate with the units *μ*mol cm^-1^ h^-1^ or *μ*mol g^-1^ h^-1^. The 0 h and 1 h samples were processed for the low nutrient treatment and the 0 h and 4 h samples were processed for the high nutrient treatment as both of these provided a measurable depletion rate for all macronutrients. The total root respiration rate was divided by the total root length to give the specific root respiration rate (nmol CO_2_ m^-1^ s^-1^). The specific root length (cm g^-1^) was calculated by dividing the total root length by the root dry weight. Broad-sense heritability (*h*2) was calculated using the equation:

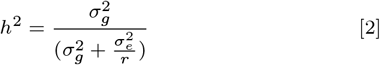

where 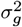 and 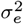 are the genetic and residual variances, respectively, and r is the number of experimental replications (61). Due to missing samples from the seven individual replicate runs, we used the average number of replications per line, which was 6.

## ACKNOWLEDGMENTS

The authors would like to acknowledge the USDA/National Institute of Food and Agriculture (Grant reference 2017-67007-25948) and Noble Research Institute, LLC for financial support. In addition, the authors would like to thank Yuhong Tang, Guifeng Li, Nick Krom, Taegun Kwon, Levi Hartman, Bonnie Watson and Michael Cloyde for technical assistance. YT and GL assisted with RNA-sequencing, NK assisted with RNA-seq analysis, TK assisted with qPCR, LH assisted with shoot imaging and shoot grinding, MC for shoot and root grinding and BW for operation of the Elemental Analyzer instrument for Dumas. The authors thank Elison Blancaflor for reviewing the manuscript for scientific soundness and providing helpful comments.

